# Maintenance of colour memoranda in activity-quiescent working memory states: Evidence from impulse perturbation

**DOI:** 10.1101/2023.07.03.547526

**Authors:** Güven Kandemir, Sophia A. Wilhelm, Nikolai Axmacher, Elkan G. Akyürek

## Abstract

The neural mechanisms underlying working memory maintenance pose a challenge for investigation, as sustained neural activity may not always be observable. To address this, the method of impulse perturbation has been employed to examine memorized information during activity-quiescent periods. However, this approach has mainly focused on spatially localized or referenced stimuli, leaving it unclear whether non-spatial memoranda share similar neural maintenance mechanisms. This study aimed to fill this gap by applying the impulse perturbation method to working memory for colours, which are inherently non-spatial stimuli. EEG data from 30 participants performing a delayed match-to-sample task were analysed, with one of the presented items being retro-cued as task-relevant. Our findings indicate that both cued and uncued colours could be decoded from impulse-evoked activity, in contrast to previous reports on working memory for orientation gratings. Additionally, we explored colour decoding from ongoing oscillations in the alpha band and discovered that cued items could be decoded, potentially influenced by attention, whereas uncued items could not. These results suggest subtle differences between the representation of colours and stimuli with spatial properties. However, they also demonstrate that both types of information can be accessed through visual impulse perturbation, regardless of their specific neural states.

## Introduction

Working memory is a system of components that enables the maintenance of information in an accessible state in the absence of sensory stimulation (Cowan, 2017). The neural basis of working memory has been of increasing interest in recent years. One particularly striking outcome has been that although working memory maintenance has been traditionally associated with sustained neural activity during the memory delay period (Kojima & Goldman-Rakic, 1982; Goldman-Rakic,1995; Constantinidis et al., 2018), a number of studies failed to observe such sustained activity (LaRocque, et al., 2013; Lewis-Peacock, et al., 2012; Shafi et al., 2007; Watanabe & Funahashi, 2014). It was proposed that during this period without an observable neural correlate, memoranda could be retained in activity-silent states (Stokes, 2015; Wolff et al., 2017), which may rely on synaptic plasticity facilitated by elevated post-excitatory calcium and neurotransmitter levels (Buonomano & Maass, 2009; Barak & Tsodyks, 2014; Kozachkov et al., 2022; Lundqvist et al., 2018; Mongillo et al., 2008; Miller et al., 2018).

Wolff and colleagues have previously shown that the presentation of a high-contrast, standardized, but task-irrelevant stimulus during the memory delay may allow decoding of such activity-silent, or at least activity-quiescent, memory items that could not be detected from raw ongoing EEG (Wolff et al., 2017). While it is not yet exactly clear how this so-called impulse signal reveals the memory trace at a physiological level, Wolff and colleagues explained this using the analogy of sonar: The impulse allows measuring a ‘hidden’ state by attributing differences in the response to a stable stimulus to underlying differences in the network (Wolff et al., 2017). In our case, the sensory processing of an impulse signal is thought to perturb the initially ‘hidden’ memory network, generating activity that can be measured with EEG, from which the state of this network can be inferred (Stokes, 2015; Wolff et al., 2015; 2017; 2020a; 2020b). Studies using the impulse perturbation approach have reported successful results for memories of orientations (Wolff et al., 2015; 2017; 2020a; 2020b), numerosity (Fornaciai & Park, 2020), and auditory tone frequencies and sequences (Fan et al., 2021; Fan & Luo, 2022; Wolff et al., 2020b).

Although impulse perturbation has thus been used with different stimuli, it is yet uncertain whether the representational patterns observed to date will hold universally for different kinds of content that are maintained in working memory. In particular, one crucial similarity between previously tested memoranda is that they allow a transformation of the task relevant information into spatial coordinates (which may also aid memory). For example, each member of a set of orientations can be represented as different points on the edge of an imaginary circle around a fixed point on a plane surface. In a similar fashion, higher or lower numerosity and tone frequency can be easily converted to different elevation levels laid out on a similar surface (e.g., a high tone may be visualized high on the vertical axis). This possibility of representing information with spatial position leaves open the question of whether the neural signature of the maintenance of this kind of information also applies to that of non-spatial stimulus attributes.

The reliance on spatial properties in previous studies also brings confounding risks with it. For example, it might involve the deployment of spatial attention, which might affect the alpha band of the EEG in particular (Worden et al., 2000). Bae and Luck (2018) investigated the contribution of spatial attention to the decoding of orientation items in working memory. They found that while alpha band activity only conveyed information about the (attended) location of their stimuli (but see also Barbosa et al., 2021), and while decoding of ongoing EEG during the delay period mainly provided significant information about the item-specific orientation, it also reflected its location. Furthermore, differences that exist between stimuli or conditions in terms of spatial attention come with the risk that (voluntary or involuntary) eye movements may follow suit. Neural data can be confounded by correlating eye movements and gaze fixations, especially under active viewing conditions (Nishimoto et al., 2017; Thielen et al., 2018; 2019). Activity in the brain, particularly in earlier visual regions, may reflect viewpoint-specific, retinotopically-organized information that will vary considerably when the eyes move around. Such activity cannot be easily discerned from other aspects of sensory and cognitive processing, including memory maintenance (Mostert et al., 2018; Quax, et al., 2019).

Considering the theoretical limitations and potential confounding issues with regard to the exclusive reliance on spatial properties in previous impulse-based experiments on working memory maintenance, we set out in the current study to overcome these by using intrinsically non-spatial stimuli, namely colours. Colours can be decoded successfully from fMRI (Brouwer & Heeger, 2009; Persichetti et al., 2015), MEG (Hermann, et al., 2022; Rosenthal et al., 2021; Sandhaeger et al., 2019), and EEG data (Bocincova & Johnson, 2019; Hajonides et al., 2021; Sandhaeger et al., 2019). Furthermore, the colour-space is similar to orientation space with regard to parametric differences between distinct equiluminant colours (Hajonides et al., 2021). Thus, colours seem a suitable non-spatial substitute to extend earlier orientation-based pinging studies (Wolff et al, 2015; 2017; 2020a; 2020b). Apart from thus changing the memory items, we also presented them serially, at the same location, rather than lateralized, as was originally done, and rotated the colour wheel that served as the response probe on each trial, thus removing all spatial aspects from the original task.

We conducted a set of pre-registered analyses, based on those reported in the original paper by Wolff and colleagues (2017), and added decoding analyses of alpha band activity. Our results show that trial-specific colour information could be successfully predicted from the activity evoked by the visual impulse, similar to orientation gratings in earlier studies (Wolff, 2015; 2017; 2020a; 2020b). In addition to the task-relevant cued colour, dynamic impulse-driven activity also revealed the uncued colour. Conversely, while alpha power decoding yielded a sustained trace of the cued memory item, this was not the case for the uncued colour. These findings extend previous work on orientation decoding, and suggest that different features might elicit (slightly) different maintenance mechanisms. The present outcomes also highlight that impulse-driven decoding can reveal memoranda in distinct memory states, independent of the allocation of spatial attention.

## Method

### Participants

Thirty volunteers (23 female, M_age_ = 24.2, Range_age_ = 20-36) that were recruited via social media adverts participated in this study in return for monetary rewards. Participant selection relied on the successful completion of a pre-screening test, which was a shortened version of the main experiment (288 trials). The preselection cut-off criterion was ≤30° of error in at least 70 % of trials. None of the volunteers were eliminated by the pre-screening. The sample size was based on earlier studies with similar designs (e.g., Wolff et al, 2017). All participants were informed about the experimental procedures as well as the data sharing procedures, and written consent was obtained. The study was conducted in accordance with the Declaration of Helsinki (2008), and it was approved by the Ethical Committee of the Behavioural and Social Sciences Faculty of the University of Groningen (Study ID = PSY-1920-S-0385).

### Apparatus and Stimuli

The experiment took place in a well-lit chamber where participants were seated 60 cm away from a 17” Samsung 797DF CRT monitor. The refresh rate was set to 100 Hz and the resolution was 1024 by 768 pixels. All stimuli were created and presented with the freely available Psychtoolbox 3 extension for Matlab (Brainard, 1997; Kleiner, et al., 2007).

The memory items and the probe consisted of coloured disks with a visual angle of 6.69°, which were presented in the centre of the screen. Their colours were randomly drawn from RGB conversions of 48 equiluminant colours equally distanced on the CIELAB colour-wheel, which were extracted from the freely available Matlab extension MemToolbox (Suchow et al., 2013). A grey background (RGB = 128, 128, 128) was maintained throughout the experiment. A black fixation dot with a white outline (0.25° of visual angle) was displayed in the centre of the screen at all times except during the presentation of the cue. The cue was a number, “1” or “2”, indicating the serial position of the task-relevant (“cued”) item in that trial, presented in Arial font in the centre of the screen (0.5° visual angle). The impulse was a large white disk, displayed in the centre of the screen with a visual angle of 13.38°. The response screen contained the probe in the centre of the screen and a colour-wheel surrounding the probe, which had a diameter of 10.05° and a width of 0.55° visual angle. A white line reaching from the centre to the edge of the colour circle indicated the momentary probe colour on the colour-wheel. In each trial, the colour-wheel was randomly rotated and a random colour was assigned to the probe disk. Responses were collected with an Xbox controller. Following a response, a happy or a sad smiley face was presented at the centre in Arial font, indicating accuracy (i.e., whether the absolute error was less or more than 30°).

### Procedure

The experiment consisted of 1536 trials, which were completed in four consecutive sessions that were separated by breaks. Participants determined the duration of the breaks. In each session participants completed 24 blocks, and after each block an average score per block was presented as feedback. Participants could start each block by pressing the SPACE bar, after which trials continued automatically until all the trials in the block were completed. At the beginning of each block, a “Get Ready” warning was presented first, after which the trials commenced.

Each trial started with the presentation of a fixation dot, which was shown for 700 ms on a grey background. This was followed by the serial presentation of two colours for 200 ms duration, each followed by a 900 ms delay. The cue was presented next for 200 ms, and a delay of 900 ms followed it. Next, the impulse signal was presented for 100 ms, and a consecutive delay was on display for 500 ms. The response screen was displayed next and stayed on the screen until a response was submitted. When the response screen was on display, participants could move the left stick on the controller to rotate the black bar presented on the probe, and change the colour of the probe circle. Once the desired colour was selected, the response was submitted by pressing X on the controller. The response was followed by a delay of 150 ms after which feedback was presented for 300 ms. The next trial began automatically after a random jitter with a range of 500 to 800 ms.

### EEG Acquisition and Pre-processing

The EEG was recorded with Brainvision Recorder software, and a TMSI Refa 8-64/72 amplifier using 62 Ag/AgCl sintered electrodes, which were placed according to the international 10-20 system. The data were recorded in reference to the average of all electrodes at a sampling rate of 1000 Hz. The ground electrode was placed on the sternum, and eye movements were tracked via bipolar electrooculography with vertical electrodes above and below the left eye, and two horizontal electrodes on the ipsilateral sides of both eyes. The resistance at all electrodes was kept below 7 kΩ throughout the experiment.

Filtering and preprocessing were handled via the Matlab extensions Fieldtrip (Oostenveld et al., 2011) and EEGLAB (Delorme & Makeig, 2004). The data were re-referenced offline to the average of both mastoids. For the multivariate analyses on voltage values, the data were downsampled to 500 Hz, and filtered at 0.1 Hz high-pass and 40 Hz low-pass. Alpha power amplitudes were acquired by bandpass filtering the EEG signal with a 8 Hz high-pass and 12 Hz low-pass, by applying the Matlab function presented below to the data on each channel:

abs(hilbert(eegfilt(data, sample_rate, low_pass, high_pass))),

where low_pass and high_pass corresponded to the 8 and 12 Hz filters applied to our EEG data, and sample_rate corresponded to the 500 Hz sampling rate. The filter output was Hilbert transformed and the absolute of the product was calculated to get the real values of the transformation.

The voltage data and alpha power amplitudes were separately epoched to the onset of the memory items, the cue, and the impulse, covering a range starting from -150 ms relative to their onset until the onset of the next stimulus, thus forming distinct epochs for item 1, item 2, cue and impulse. Semi-automatic artefact rejection was completed by marking trials with high voltage variations and then inspecting all trials visually for channel drifts, muscle and eye artefacts. Drifting channels were interpolated using the spherical head model, whereas epochs with other artefacts and blinks were excluded from the analyses. In total 13.24% of the Item 1 epochs, 10.7% of the Item 2 epochs, 10.76% of the Cue epochs, and 9.99% of the impulse epochs were excluded.

### Multivariate Analyses

Unless stated otherwise, all analyses were pre-registered at https://osf.io/uvxe7/. The decoding analyses were restricted to the 17 posterior channels (P7, P5, P3, P1, Pz, P2, P4, P6, P8, PO7, PO3, POz, PO4, PO8, O1, Oz and O2), replicating earlier studies that investigated visual working memory by means of impulse perturbation (Wolff et al., 2017; 2020a; 2020b).

### Analysing the time window of interest

The primary analysis aimed to investigate the accuracy of trial-specific colour decoding, which was calculated using the data within a time window of interest. This time window of interest covered 100-400 ms relative to the presentation of a stimulus (e.g., memory item or impulse). This window was based on earlier studies, in which it was applied to capture the bulk of the EEG response to the eliciting stimulus, and in particular the dynamic response to the impulse (Wolff et al., 2015; 2017). First, the data were baselined by subtracting the average activity within the aforementioned period, which was done separately for each trial and electrode. Next, the data were downsampled by calculating the moving average over a 10 ms window. These downsampled values were then pooled over all 17 posterior electrodes, which yielded a single spatio-temporal pattern for each trial.

The trials were assigned to the closest one of 16 equidistant bins that covered the pre-determined colour-space, and were re-labelled with the centre value of that bin. This was repeated three times to cover the pre-determined colour-space (from 0° to 337.5°, 7.5° to 345°, and 15° to 352°, each in steps of 22.5°), thus providing 16 different colour conditions in each of the three runs. Next, the trials were partitioned into 8 folds with seven folds serving as the training set. Within the training set, the number of trials in each colour condition were equalized by subsampling the data. Subsequently, the data within each colour condition were averaged, forming 16 condition-specific spatio-temporal patterns. These spatio-temporal patterns of colour conditions were then convolved with a half cosine basis set raised to the 15^th^ power to reduce noise and pool information across similar colours. The similarity between test trials and the averaged training data was quantified in Mahalanobis distances (De Maesschalck et al., 2000), yielding 16 distance values for each test trial. The covariance matrix was estimated from the entire training set by using a shrinkage estimator (Ledoit & Wolf, 2004). The distance values were mean-centred and sign-reversed, so that positive values indicated higher similarity. Finally, the values were convolved with a cosine-similarity function of the colour space (i.e., on the colour wheel). The product was the trial-specific decoding accuracy. The procedure was repeated 100 times with random folds and random sub-sampling, in order to avoid sampling biases. The reported decoding accuracy for each participant was calculated by averaging all the products of all repetitions and all trials to get a single value per participant.

### Time-course analyses

The time-course of the memory-related dynamic signal was investigated by sliding a 100 ms time-window across the epoch. In each step, data within the window were baselined by subtracting the mean activity within, and then the residual activity was downsampled to 100 Hz. The data were then pooled over electrode space, yielding the spatio-temporal pattern at that time point. The time-course analysis was highly similar to the analysis of the time window of interest. The trials were first re-labelled in accordance to the colour-space. The trials were divided into 8 folds with stratified sampling so that all conditions had an approximately equal representation in each fold. The training was set formed by 7 folds and the trials in these folds were distributed across 16 bins. The number of trials in each bin was equalized with random subsampling in order to avoid bias, and the data in each bin was averaged to form temporal patterns for each colour bin at each time point. The measure of similarity between the test trials and the 16 averaged colour patterns was calculated at each time point in Mahalanobis distances. A shrinkage estimator was used to estimate the covariance matrix from the training set (Ledoit & Wolf, 2004). The distance measures at each time point were reverse-signed and mean-centred, and then scaled by cosine convolution of the modelled colour space. The output was the decoding accuracy at each time point. The procedure was performed 3 times, once for each colour-space, and repeated 100 times to avoid sampling bias. The output was averaged over colour spaces, repetitions and trials for each participant.

Post-hoc, we conducted another time-course analysis on (absolute change in) alpha power, which was not included in the preregistration. The analysis was identical to the above, but applied to filtered 8-12 Hz EEG data, which was baselined over -200 to 0 ms relative to cue and impulse onset. The data were also downsampled to 125 Hz to save computational time. Finally, we also conducted the same time-course analysis on the EOG data to ensure the absence of any spatial correlation.

### Representational similarity analyses

Representational similarity analyses (RSA) (Kriegskorte et al., 2008) was used to parametrically assess the relationship between colours during memory encoding and during the memory delay following the impulse. For the RSA, the EEG data from the same pre-determined 17 posterior electrodes within the time window (100 to 400 ms relative to a stimulus or an impulse signal onset) were prepared as in the decoding analysis of the time window of interest. The trials were re-labelled according to the first colour-space (0° – 337.5° on the colour wheel) and grouped into 16 bins. The number of trials in each bin were equalized and 16 spatio-temporal patterns were generated by averaging trials in each bin. The pairwise differences between all bins were calculated in Mahalanobis distances to form the representational dissimilarity matrix (RDM). The covariance matrix was estimated from the training set with the use of a shrinkage estimator (Ledoit & Wolf, 2004). The entire procedure was repeated 100 times to account for selection biases and the final RDM was averaged over all repetitions for each participant.

The RDM was tested by the linear regression of two models. The first model covered the circular nature of 16 colour bins centred according to the first colour space. In other words, this model specifically assessed the hue differences in terms of angular space on the colour wheel. The second model was based on the behavioural output, which showed that the responses clustered around three primary colours. The second model thus assessed if the differences in neural space for the 16 colour bins could be explained by the differences between the three frequently reported discrete colours (green-blue, red and yellow). Both models were first converted to z-scores. Next, the RDM of each participant was regressed against each model separately. The diagonal segment for the RDM and models was excluded from analyses. The standardized slopes (beta values) were taken as an indication of the fit. The mean beta values were contrasted against zero by using a group permutation test.

### Statistical assessment

Trial-wise decoding accuracy was averaged for each participant, yielding a single output (at each time point) per participant. The average decoding accuracy over all participants was contrasted against a null distribution (one-sided). In order to form this null distribution, the sign of the mean decoding accuracy (per participant) was flipped 100,000 times with 50 % probability, and the resultant mean was taken. The proportion of the null distribution larger than the observed mean accuracy was calculated as the *p* value, which was labelled as significant if it was smaller than 5% (*p* < 0.05). For the time-course analysis, an additional cluster correction was applied to control for multiple comparisons, where the cut-off was set to 0.05 (*p* < 0.05). Confidence intervals were built by bootstrapping the mean (n_perm_ = 100,000). When decoding accuracies for two conditions were contrasted, the difference of these two conditions was calculated for each participant, and this group vector was tested against zero by using the same sort of permutation test as explained above. The RSA was similarly assessed by means of a group permutation test (n_perm_ = 100,000), which was applied on the beta values of the participants, reflecting the model fit to the pairwise differences reported in the RDMs.

### Data availability

The data and analysis scripts of this study are publicly available at https://osf.io/bxmt8.

## Results

### Behavioural Results

The overall behavioural performance in the experiment was good; the mean error was 18.3° with a standard deviation of 32.5° (Figure 2A), relative to the centre of the bin to which the cued item belonged (adjusted error). Figure 2B presents the behavioural reports of the cued item, with the uniform colour bins overlaid. This plot reveals that despite the uniform distribution of the colours presented during the experiment, the reported colours seemed clustered around the primary hues, supporting earlier evidence that memory performance varies for different colours (Bae et al., 2014).

**Figure 1.**
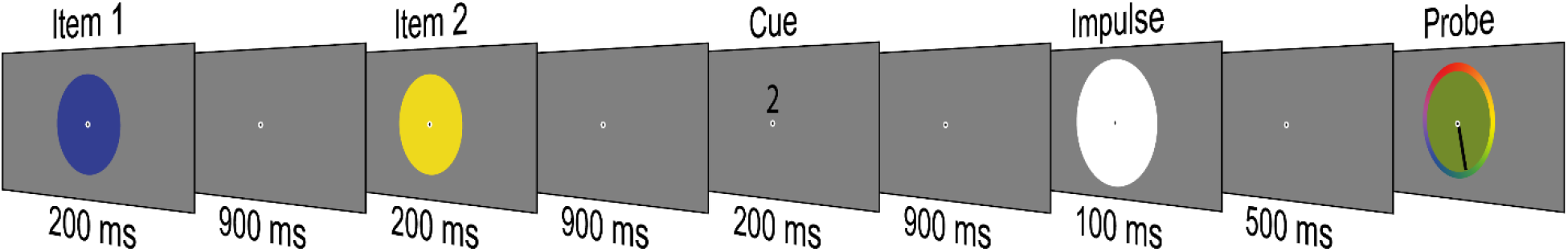
Overview of a single trial in the experiment. In each trial, the colours of two sequentially presented disks would be memorized. A retro-cue then indicated the temporal position of the task-relevant memory item. Finally, the randomly-oriented probe was rotated by the participants with a joystick to report the task-relevant colour. The numbers below the stimulus displays reflect their duration in ms.

**Figure 2.**
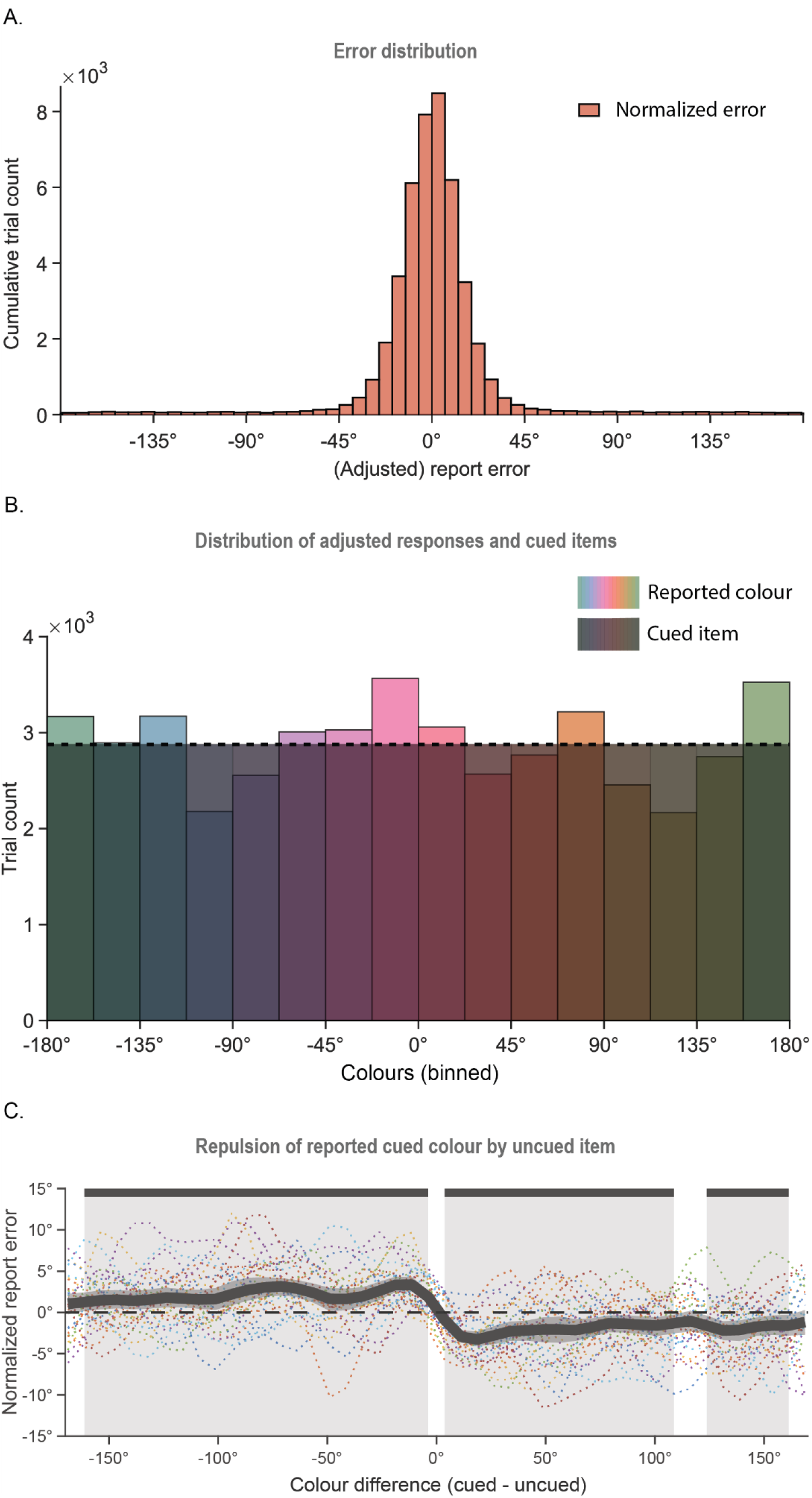
Behavioural plots. **(A)** Histogram of report errors relative to the task-relevant memory item. **(B)** Distribution of the cued colour bins (shaded, dashed line) and the reported colour bins, as a function of the angular values of the colour-space. **(C)** Normalized error as a function of the difference between the cued and the uncued item. The moving mean of the report error is calculated for a 22.5° wide bin in steps of 7.5°. The shaded area and the solid bars at the top mark the colour differences for which the adjusted error differed from 0°, according to the cluster-corrected permutation test (*p* < 0.05).

The influence of the uncued memory item on the report of the cued item was also investigated. Since only the cued item was reported in each trial, the presence of the uncued item was assessed as the effect of the similarity between the cued and the uncued colour on the degree of error. To take into consideration possible individual differences in colour perception (Emery & Webster, 2019), errors were first median-normalized within each colour bin. The normalized error values were then binned again as a function of the difference between the task-relevant cued item and the task-irrelevant uncued item (binwidth = 22.5°, moving window in steps of 7.5°), and the mean error was calculated within each bin. A permutation test was applied with cluster correction to assess the deviation of the mean from zero at each unit of angular difference between the items. The report error was significantly different from zero for three difference ranges (Figure 2C, M_Adjusted error_ ≠ 0, for cued – uncued difference, ranging from -161° to -3.75°, *p* < 0.001, from 3.75° to 116°, *p* < 0.001, and from 124° to 161°, *p* = 0.024). The cued item report errors deviated away from the uncued item, reflecting a repulsion away from the task-irrelevant colour.

### Decoding the time window of interest

Both the trial-specific colour of item 1 (Figure 3A, left, red, *p* < 0.001, one-tailed), and of item 2 (Figure 3A, left, blue, *p* < 0.001, one-tailed), could be successfully decoded after presentation. Interestingly, item 1 was also decodable within the critical period following the presentation of item 2 (Figure 3A, left, item 1 (2), red, *p* < 0.001, one-tailed). Decoding of item 1 after the onset of item 2 was nevertheless significantly lower than stimulus-driven decoding (difference _item 1 – item1 (2)_, *p* < 0.001, one-tailed). The associated tuning curves (Figure 3A, right) reflected a parametric relationship between colours, in line with earlier reports (Hajonides et al., 2021).

**Figure 3.**
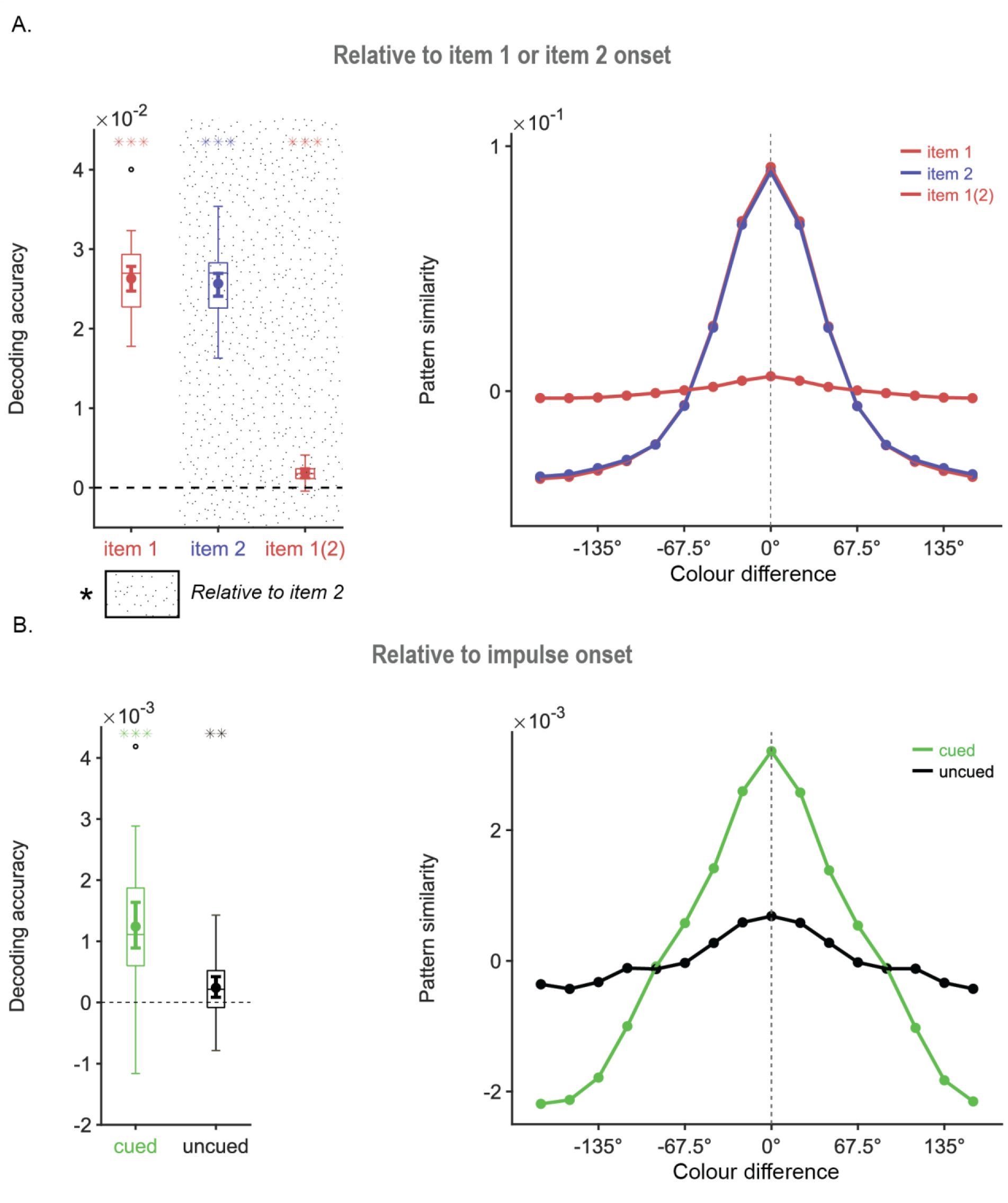
Boxplots and tuning curves showing decoding accuracy and pattern similarity (in arbitrary units) for memory items within the 100-400 ms time window of interest relative to the onset of item 1 and 2 presentation (A), as well as impulse onset (B). **(A)** Item 1 (red) could be successfully decoded after its onset, and after onset of item 2 (“item 1(2)”). Item 2 (blue) was also successfully decoded after its onset. The pattern similarity plots (A, right) show a circular, parametric relationship between colours, reflected by the gradual change in pattern similarity as a function of the colour difference for item 1 and item 2. **(B)** Cued (green) and uncued (grey) items were both successfully decoded from the time window of interest, after impulse onset. The associated pattern similarity plots show that the parametric relationship between memory items was preserved (B, right). The boxes border the 25^th^ and 75^th^ percentiles, with the whiskers around the box stretching to 1.5 interquartile range. Asterisks indicate beta values that were significantly above zero (*, *p* < 0.05; **, *p* < 0.01; ***, *p* < 0.001).

After impulse presentation, not only the task-relevant, cued colour was decodable (Figure 3B, left, green, *p* < 0.001, one-tailed), but also the task-irrelevant uncued item (Figure 3B, left, grey, *p* = 0.008, one-tailed), in contrast to earlier studies that used orientation stimuli (e.g., Wolff et al., 2017). Although both cued and uncued memories could be traced, there was a clear difference in the strength of their representations (difference _cued – uncued_, *p* < 0.001, one-tailed). Both the cued and uncued item showed parametric pattern similarity, as expected (Figure 3B, right).

### Time-course decoding

In addition to classic time-course decoding of colours, we additionally applied this analysis to alpha power. Motivating this addition was the recent suggestion that ongoing activity in the alpha band could reflect memory content in a sustained fashion, which questions the need to use the impulse perturbation method (Barbosa et al., 2021). The analyses were focused on the cue and impulse epochs, where the selection between task-relevant (cued) and task-irrelevant (uncued) items had been made. Furthermore, we were able to confirm that eye movements, attentionally driven or otherwise, did not affect decoding of the memory items by applying the same analysis to the EOG data (Supplementary Information).

The memory items could not be decoded from the voltage data following the presentation of the cue (Figure 4A, B). Conversely, both the cued and the uncued items were successfully decoded from alpha power in an earlier phase after the presentation of the cue (Figure 4C, green, 396 ms to 588 ms, *p _cued_* = 0.015, two-tailed, corrected; Figure 4C, black, 364 ms to 516 ms, *p _uncued_* = 0.014, two-tailed, corrected). Subsequently, in a later phase, only the task-relevant, cued colour was decodable in the alpha band without interruption for the remainder of the epoch until the onset of the impulse signal (Figure 4C, green, 700 ms to 1050 ms, *p _cued_* < 0.001, two-tailed, corrected). The difference in decoding accuracy between cued and uncued items was also statistically significant for some time within this period, from 892 ms to 1028 ms relative to impulse onset (*p _cued - uncued_* = 0.037, one-tailed, corrected). Across these phases, parametric pattern similarity was also apparent for both items (Figure 4D).

**Figure 4.**
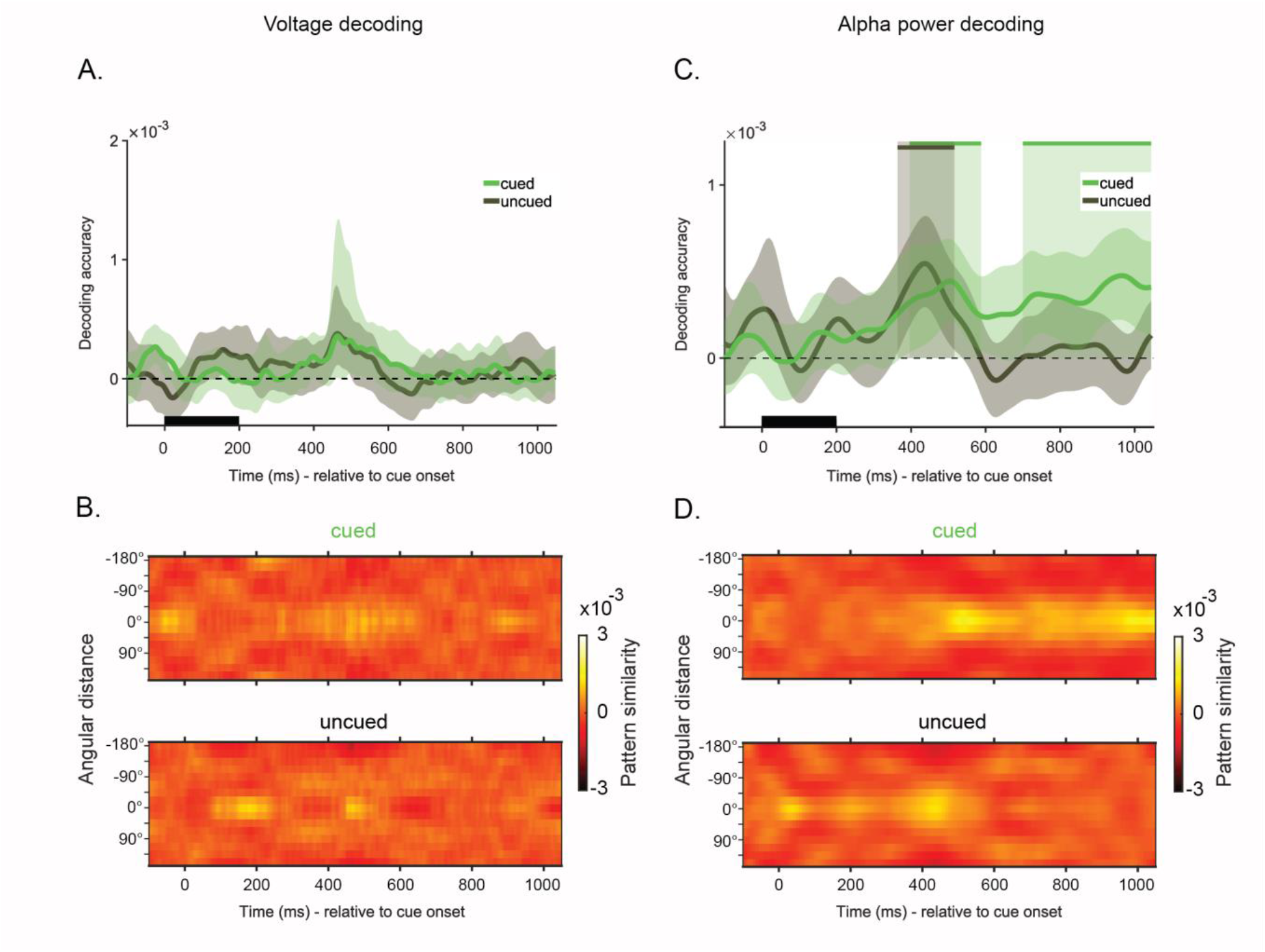
Time-course decoding of the cued (green) and the uncued (black) items, relative to the onset of the cue from the raw voltages **(A-B)**, and from alpha power **(C-D)**. **(A-C)** The black rectangular bar marks the presentation of the cue. Solid lines show the mean decoding accuracy (A.U.) over all trials and participants as a function of time. The shaded area around the mean marks the 95 % CI. Solid bars at the top and the shaded zones indicate statistically significant decoding periods (*p* < 0.05, one-sided). **(B-D)** Pattern similarity matrices for cued and uncued items show reverse-signed, mean-centred Mahalanobis distances between the target item and all other possible memory items, averaged over trials as a function of time.

At impulse presentation, voltage decoding revealed both the cued (Figure 5A, green, 28 ms to 514 ms, *p _cued_* < 0.001, two-tailed, corrected), and the uncued item (Figure 5A, black, 164 ms to 236 ms, *p _uncued_* = 0.036, two-tailed, corrected). The difference between the states of these two items was reflected by a significant difference in decoding accuracy (118 ms to 356 ms, *p _cued - uncued_* = 0.005, one-tailed, corrected). Pattern similarity reflected this difference also, but was qualitatively similar for both items (Figure 5B). These results confirmed the outcomes of the analysis of the time window of interest reported above. The cued item was also decodable from alpha power (Figure 5C, green, 68 ms to 550 ms, *p* < 0.001, two-tailed, corrected), but the uncued item was not (difference; 92 ms to 548 ms, *p _cued - uncued_* < 0.001, one-tailed, corrected). The pattern similarity matrix of the cued item showed high similarity near the tested values, with a steep drop-off further away (Figure 5D). These results suggest that while alpha power may reflect the sustained maintenance of, or the attention allocated to, the task-relevant memory item, the task-irrelevant item only emerged from the activity evoked by the impulse. Given that the uncued item also biased the eventual behavioural response, the present findings provide evidence that the impulse signal can reveal memory items retained in different activity states (Stokes, 2015; Stokes et al., 2020).

**Figure 5.**
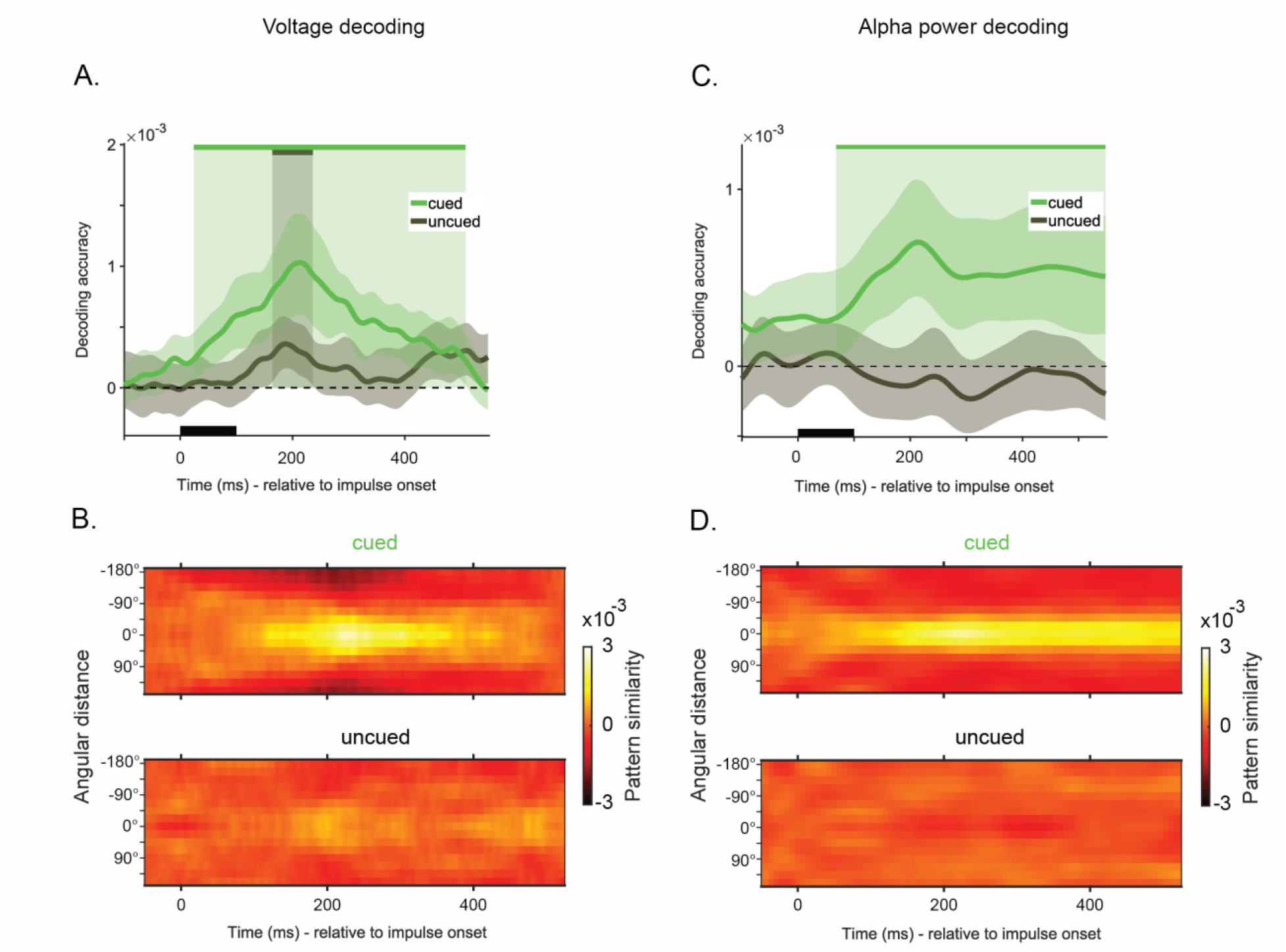
Time-course decoding of the cued (green) and the uncued (black) items, relative to impulse onset from the raw voltages **(A-B)**, and from alpha power **(C-D)**. **(A-C)** The black rectangular bar marks the presentation of the impulse. Solid lines show the mean decoding accuracy (A.U.) over all trials and participants as a function of time. The shaded area around the mean marks the 95 % CI. Solid bars at the top and the shaded zones indicate statistically significant decoding periods (*p* < 0.05, one-sided). **(B-D)** Pattern similarity matrices for cued and uncued items show reverse-signed, mean-centred Mahalanobis distances between the target item and all other possible memory items, averaged over trials as a function of time.

### RSA of Colour Representations

In the RSA, two models were tested to explain pairwise differences between colour bins. The first model was based on a uniform colour space that reflected the angular differences between 16 colour bins on the colour wheel (Figure 6A, left). The second model was based on the behavioural reports (Figure 6A, right), in which errors were found to be more frequent around three primary colours (Figure 2B). In the model, these three colours were thus used to explain the variation in the data. The average RDMs presented in Figure 6B reflect the pairwise differences between the 16 colour bins within the time window of 100 to 400 ms, relative to the onset of the memory items (left), and relative to impulse onset (right).

**Figure 6.**
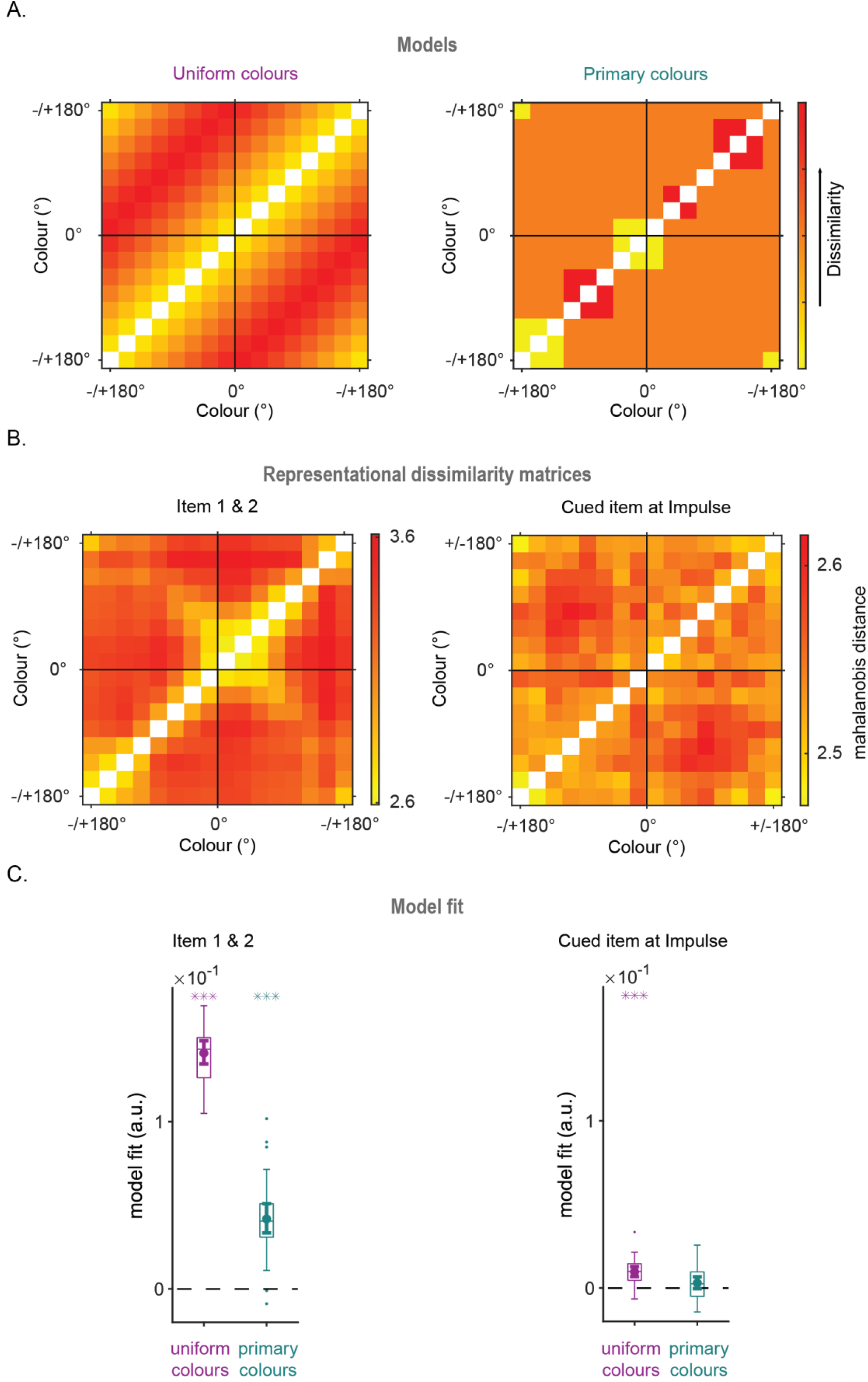
Representational similarity analysis of the average of item 1 and 2 (left) and the cued item at impulse (right). **(A)** Model for a uniform, continuous colour space for 16 colour bins (left), and model for a discrete colour space based on primary colours extracted from behavioural errors (right). **(B)** The representational dissimilarity matrices (RDM) for item presentation (left) and impulse (right) epochs. **(C)** Boxplots showing the mean fit (beta values) following the linear regression of individual RDMs on the models for each participant. The dot in the centre marks the mean standardized slope (beta), and the whiskers on both ends of the mean indicate the 95% CI. The boxes border the 25^th^ and 75^th^ percentiles, with the whiskers around the box stretching to 1.5 interquartile range. Asterisks indicate beta values that were significantly above zero (*, *p* < 0.05; **, *p* < 0.01; ***, *p* < 0.001).

Both models fit the data during stimulus encoding (Figure 6C, left, *p _Uniform colours_* < 0.001, one-tailed; *p _Primary colours_* < 0.001, one-tailed). Thus, the results provided evidence for a parametric relationship between colour representations during sensory encoding, as well as a degree of primary-colour categorization. The uniform colour space model also fit the neural response associated with the cued item that was evoked by the impulse (Figure 6C, right, *p _Uniform colours_* < 0.001, one-tailed), in line with earlier reports (Hajonides et al, 2021). Conversely, although there was a trend, the primary colours model failed to reach statistical significance (Figure 6C, right, *p _Primary colours_* = 0.054, one-tailed).

## Discussion

We aimed to decode colours maintained in working memory by means of visual impulse perturbation. In a pre-registered experiment, we tested the maintenance of a retro-actively cued target item, and of the item that was not cued, in a delayed match-to-sample task. To further avoid possible spatial confounds that might have affected previous impulse perturbation experiments, we presented our two memory items serially in the centre of the screen, and randomized the appearance of the colour circle shown at the probe on each trial. We were able to decode stimulus identity from the recorded EEG signal during perceptual encoding, replicating earlier studies (Bocincova & Johnson, 2019; Brouwer & Heeger, 2009; Hajonides et al., 2021; Hermann et al., 2022; Persichetti et al., 2015; Rosenthal et al., 2021; Sandhaeger et al., 2019). Importantly, we could also decode stimulus identity post-impulse. This result extended earlier studies that decoded the bottom-up activity induced by the sensory processing of a visual impulse signal to reveal representations of orientation gratings in working memory that were otherwise not traceable from raw EEG (e.g., Wolff et. al., 2015; 2017; 2020). The present result was the first demonstration of impulse-driven decoding of non-spatial features, specifically colour memories, thereby generalizing the findings across feature dimensions.

In contrast to previous studies (Kuo et al., 2012; LaRocque et al., 2013; Lewis-Peacock et al., 2012; Wolff et al., 2017; 2020a; 2020b), the impulse effect was not restricted to the task-relevant cued item. This suggests that the task-irrelevant uncued item was also maintained in memory, albeit to a lesser degree. One trivial reason for this might be that participants simply selected the wrong item on some trials, but there is reason to doubt this explanation. First, in alpha power decoding (discussed further below), the signal associated with the uncued item behaved completely different from the cued item, both during maintenance and following the impulse. Second, behavioural response errors on the cued item were biased by the uncued item (cf. Bae& Luck, 2017; Golomb, 2015; Huang & Sekuler, 2010), but they were biased away from it, rather than towards it—the latter would be expected when the uncued item was erroneously reported.

The emergence of the uncued item after impulse onset casts doubt on the idea that information in working memory that is no longer needed is actively purged. Memory might alternatively ‘let go of’ task-irrelevant items, for instance by no longer periodically refreshing them, such that their representations fade relatively quickly, but might nevertheless still linger for some time. This idea is compatible also with a recent model of working memory based on calcium-mediated short-term synaptic plasticity (Pals et al., 2020). The absence of task-irrelevant items in neural measures obtained in previous studies might be a consequence of the inherent weakness of their representations, which makes them harder to detect than task-relevant items in the first place. The high number of trials in the current study may have helped to overcome this difficulty. However, from the present data it cannot be excluded that the presence of the uncued item during memory maintenance might be related specifically to the metathetic nature of colour. This awaits further experimentation.

The decoding analyses showed that alpha band decoding and dynamic voltage decoding clearly reflected different states of the memoranda. After cue onset, item selection following the processing of the cue was only observable in alpha power. Both the task-relevant and irrelevant colours could initially be decoded, followed by a sustained signal for the cued item only. With regard to the cued item, the presentation of the cue would be hypothesised to require an active process of prioritizing one item over the other, or potentially even removing the uncued item from memory storage (but see the argument above). Even theories of activity-silent working memory have so far proposed connectivity-based, synaptic storage only as a mechanism of maintenance, and updating of synaptic weights would still require neuronal firing (Kozachkov et al., 2022; Pals et al., 2020; Stokes et al., 2015; Trubutschek et al., 2019; Wolff et al., 2021). Based on the analyses of our current dataset it appears that this attentional selection happens uniquely in the alpha band, which subsequently then also holds the prioritized item in a sustained manner for the rest of the trial. With regard to the uncued item, the absence of a sustained signal both in the raw EEG and in the alpha band suggests that it was truly activity-quiescent, if not altogether silent.

Once the impulse was presented, the voltage decoding revealed both the cued and uncued item, contrary to the alpha band decoding. Additionally, the reactivation of the signal in the voltage decoding was time-limited, and returned to zero well before the presentation of the probe. This provides further evidence that the alpha band and voltage decoding track functionally different states of working memory. We speculate that the alpha band may serve to keep memoranda in an elevated state, ready for direct access when necessary (Oberauer, 2001; 2002). It seems likely that attention mediates this (Bae & Luck, 2018). Following the cue, the uncued item was apparently removed from this elevated state, but it was not altogether lost, as dynamic voltage decoding still revealed it from the EEG response to the impulse. Although the functional states of items in memory need not necessarily map directly onto corresponding neural states (Muhle-Karbe et al., 2021; Stokes et al., 2020), the task-irrelevant uncued item was clearly in a different, more silent, neural state than the task-relevant cued item in our data. The fact that we can trace these differences and differentiate them provides further support for the utility of the impulse driven decoding technique.

Finally, we found that the neural representations of colours were parametrically arranged during encoding (Brouwer & Heeger, 2009; Hajonides et al., 2021). The signal evoked by the impulse also showed a parametric arrangement of colours during the delay period. Nevertheless, behavioural responses reflected a bias in reports, as the errors were grouped around three primary colours. This observation was in line with earlier reports that not all colours are equally memorable (Bae et al, 2014). A discrete colour model derived from the behavioural output was indeed also supported during the encoding of the memory items (i.e., directly after their onset), suggesting that participants formed a categorical representation for colours even during perception or shortly thereafter. Crucially, the evidence for this discrete model was no longer reliable during memory maintenance (i.e., after the impulse), while the continuous model was still well supported. It may appear paradoxical that the categorical representation observed during encoding, and in the eventual response, was not clearly represented during memory maintenance. One possible explanation for this could be the differential impact of the activity induced by the sensory processing of the impulse signal on the different networks that might retain continuous colour representations and discrete categorical representations. One might suppose that more discrete colour groupings could be represented by a semantic network (Rosenthal, et al., 2021), whereas more low-level hue differences could be retained in earlier visual areas (Brouwer & Heeger, 2009). The latter might be more accessible to impulse perturbation.

### Conclusion

The results of our study highlighted both commonalities and potential differences between spatial and non-spatial (colour) items maintained in working memory. First, we observed that the visual impulse response contained information not only about the task-relevant, cued item, but also the task-irrelevant, uncued item, contrary to previous studies of orientation gratings. This finding casts doubt on the idea that uncued items are actively purged from memory. Second, we found evidence for a sustained signal corresponding to the cued, but not the uncued, item during the delay period. This suggests that the alpha band may uniquely trace the attended item that is in the focus of attention. Furthering the debate on the relationship between the functional role of an item and its activity state (Barbosa et al., 2021; Stokes et al., 2020; Wolff et al., 2021), the current results provided evidence for the utility of the impulse technique by revealing silent memories that were not only hidden in ongoing EEG, but also untraceable by alpha power.

## Acknowledgements

The authors are indebted to Mark Stokes and Michael Wolff for their advice in the earlier phases of this research project. This research was in part funded by an Open Research Area grant to EGA (NWO 464.18.114) and NA (DFG project number 396894956).

## Supplementary Information

To ensure that the decoding of the memory items was not confounded by (attentionally driven) eye movements, we decoded the bipolar voltage measures taken from the eye electrodes. The decoding approach was identical to that of the time course analysis of the EEG. The statistical analyses revealed no statistically significant clusters for which decoding accuracy was above zero.

**Figure S1.**
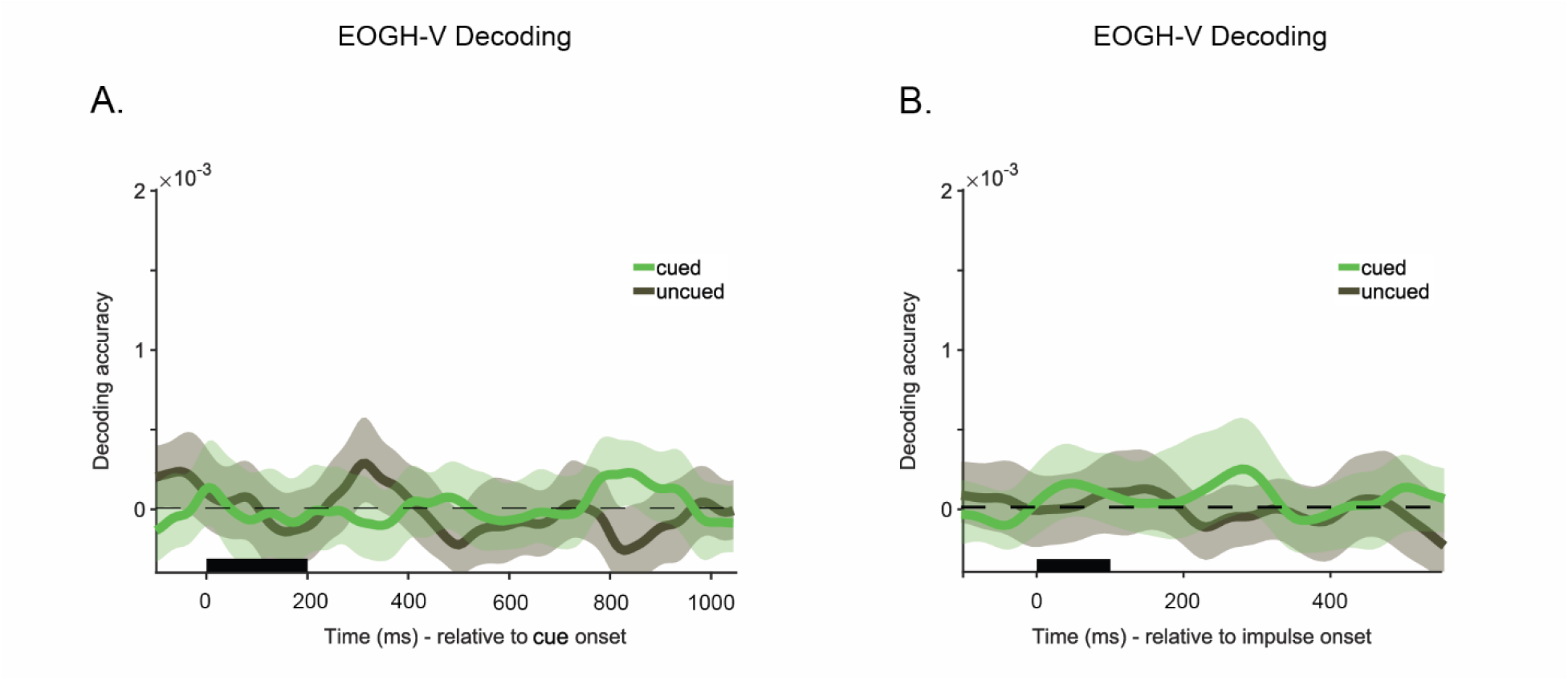
Time-course decoding of the cued (green) and the uncued (black) items, relative to cue onset (A) and impulse onset (B) from the voltage measures from the bipolar eye electrodes. The black rectangular bar marks the presentation of the cue (A) and the impulse (B). Solid lines show the mean decoding accuracy over all trials and participants as a function of time. The shaded area around the mean marks the 95 % CI.

